# Metabolic engineering of *Methanothermobacter thermautotrophicus* ΔH for recombinant acetoin production

**DOI:** 10.64898/2025.12.10.693358

**Authors:** Tina Baur, Maximilliene T. Allaart, Aaron Zipperle, Gabriela Contreras, Christian Fink, Largus T. Angenent, Bastian Molitor

## Abstract

Thermophilic methanogens of the genus *Methanothermobacter* are established biocatalysts in power-to-gas applications, converting H_2_ and CO_2_ into CH_4_ through the process of methanogenesis. Further expanding this platform for the bioproduction of value-added compounds (power-to-x) has the potential to increase the economic viability of such processes. This requires a genetic toolset that enables the controlled expression of recombinant pathways. Here, we report the fully autotrophic inducible recombinant bioproduction of acetoin from H_2_ and CO_2_ in *Methanothermobacter thermautotrophicus* ΔH. To facilitate inducible gene expression, we implemented an anhydrotetracycline (aTc)-inducible promoter system, expanding our available set of promoters. The aTc-inducible system enabled controlled expression of a codon-optimized acetoin-production operon comprising the acetolactate synthase- and acetolactate decarboxylase-encoding genes from *Streptococcus thermophilus*. Batch cultivation at 42°C demonstrated aTc-dependent acetoin formation, yielding up to 0.45 ± 0.08 mM acetoin. Fed-batch bioreactor experiments confirmed growth-coupled, recombinant acetoin production, while eliminating the non-specific acetoin accumulation that we observed during non-growth phases in batch cultivation. Continuous cultivation in a chemostat resulted in stable acetoin production rates of 1.28 ± 0.07 µmol L⁻¹ h⁻¹ at 42°C. Elevated temperatures led to reduced acetoin production, suggesting diminished activity or thermal instability of the heterologous enzymes. This study demonstrates the feasibility of value-added bioproduction in *Methanothermobacter* and establishes an inducible expression system suitable for pathway engineering in thermophilic methanogens. Together with genome-scale modeling and emerging enzyme engineering strategies, these results lay the foundation for developing robust, CH_4_-co-producing power-to-x bioprocesses with *Methanothermobacter* species.

## 1. Introduction

Methanogenic archaea (methanogens) are utilized biotechnologically as biocatalysts in power-to-gas technology, specifically for biomethanation (Pfeifer et al., 2020). In this process, renewable electric power is converted to molecular hydrogen (H_2_) *via* electrolysis of water and combined with carbon dioxide (CO_2_) from industrial off-gases for the conversion to methane (CH_4_) by the unique metabolism of methanogens, which is methanogenesis (Angenent et al., 2022; Thauer et al., 2008). Specifically, axenic cultures of thermophilic methanogens of the order Methanobacteriales (*i.e.*, *Methanothermobacter* species) have been adopted to large-scale processes with a high technological readiness level (Contreras et al., 2022; Martin et al., 2013; Pfeifer et al., 2020; Thema et al., 2019). The production of CH_4_ from surplus renewable electric power presents an opportunity to store this energy for later use in the energy system or for chemical industries with high storage capacities in various locations (Angenent et al., 2024; Wulf et al., 2020).

To improve the economics of power-to-gas technology, metabolic engineering of methanogens can be considered for the bioproduction of higher-value recombinant products (power-to-x), in addition to CH_4_ (Enzmann et al., 2018; Mühling et al., 2024). The bioproduction with thermophilic methanogens offers opportunities with distinct advantages, such as lower contamination risk, robust bioprocessing, facilitated separation of volatile compounds *via* gas stripping, and CH_4_ as the sole by-product (Contreras et al., 2022; Mühling et al., 2024; Pfeifer et al., 2020). We have previously established genetic tools for the manipulation of *Methanothermobacter thermautotrophicus* ΔH (hereafter *M. thermautotrophicus*), which provided the foundation for this endeavor (Fink et al., 2022; Fink et al., 2021; Fink et al., 2023). Recently, these genetic tools have been adapted for a closely related species, *Methanothermobacter marburgensis*, to implement markerless mutagenesis, producing isoleucine exclusively from threonine, which widens the possible strain selection for bioproduction (Klein et al., 2025).

Here, we demonstrate the bioproduction of the first recombinant product, exemplified with acetoin, from H_2_ and CO_2_ with *M. thermautotrophicus*. Acetoin has applications as a fragrance and flavor enhancer, and commercial acetoin production currently mostly relies on chemical synthesis from fossil feedstocks (Cui et al., 2022). Its bioproduction is often achieved in heterotrophic hosts utilizing carbohydrates (Cui et al., 2022; Xiao and Lu, 2014). Autotrophic bioproduction of acetoin in gas fermentation was demonstrated with *Moorella thermoacetica* (Kato et al., 2024), *Cupriavidus necator* (Windhorst and Gescher, 2019), and *Eubacterium limosum* (Shin et al., 2023), while those microbes produce soluble by-products, such as acetic acid. Autotrophic bioproduction with *M. thermautotrophicus* would allow us to integrate this process with power-to-gas technology (power-to-x), with only gaseous CH_4_ as a by-product. To achieve bioproduction in *M. thermautotrophicus*, we established an anhydrotetracycline (aTc)-inducible promoter system to augment our genetic toolbox, which had previously only contained constitutive promoters (Fink et al., 2021). We based the aTc-inducible promoter on the *tetR*-P*_tet_* system, which is widely applied in microbes, including the mesophilic methanogen *Methanosarcina acetivorans* (Guss et al., 2008). Based on this new genetic tool, we constructed a plasmid-encoded and aTc-inducible acetoin-production operon-carrying strain. This enabled the bioconversion of H_2_ and CO_2_ to acetoin with the acetolactate synthase (AlsS) and acetolactate dehydrogenase (AlsD) from the moderately thermophilic bacterium *Streptococcus thermophilus* (**Figure 1**) (Bolotin et al., 2004; Markakiou et al., 2020; Radke-Mitchell and Sandine, 1986).

**Figure 1.**
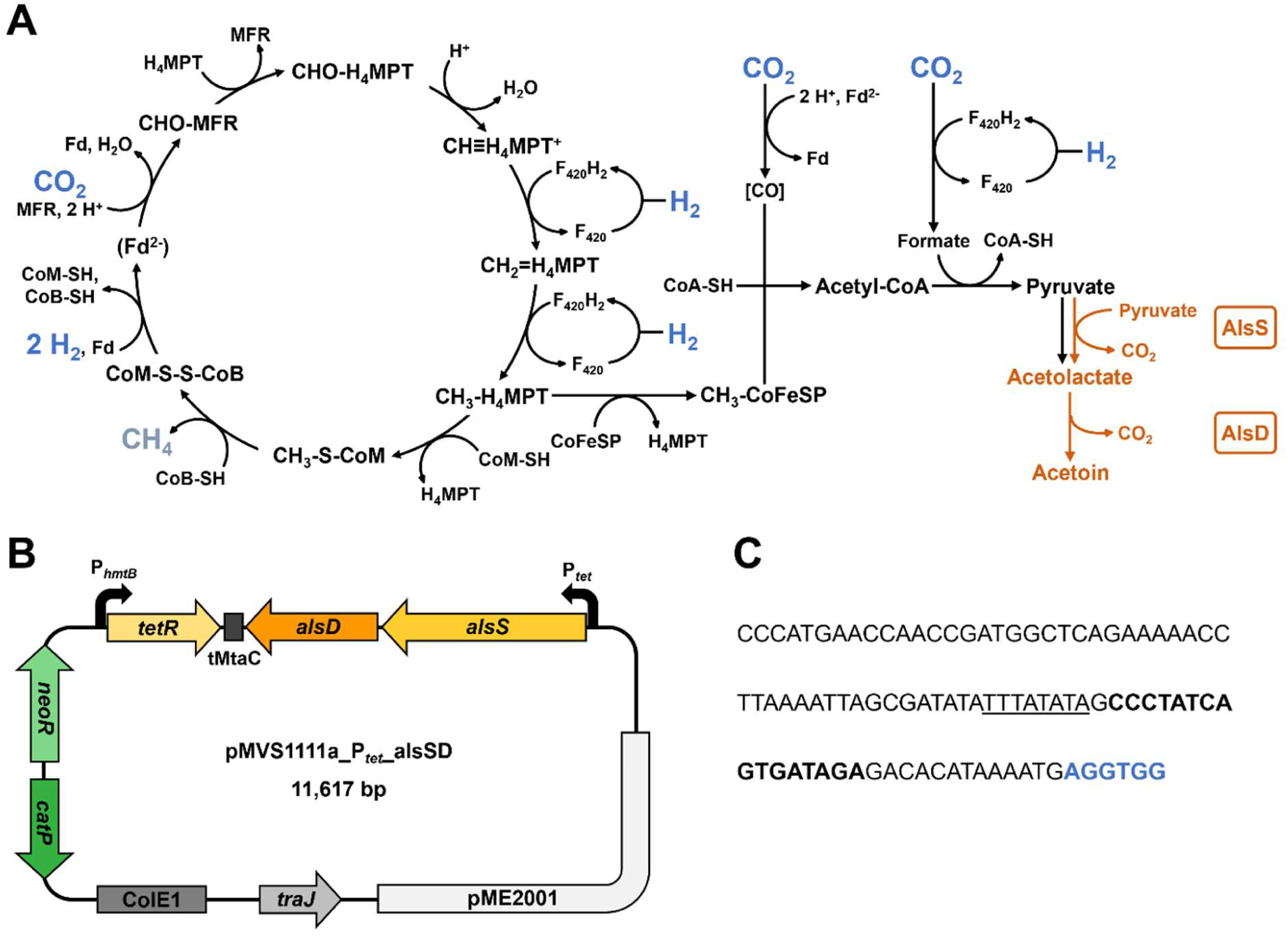
Implementation of recombinant acetoin production in *M. thermautotrophicus*. **A**, Schematic overview of the Wolfe cycle of *M. thermautotrophicus* (based on (Thauer, 2012)), and recombinant pathway for acetoin production (orange); The energetic coupling in the Wolfe cycle likely does not happen *via* free ferredoxin but within a multi-enzyme complex (Watanabe et al., 2021). **B**, Schematic overview of the acetoin production plasmid pMVS1111a_P*_tet_*_alsSD; **C**, sequence of anhydrotetracycline-inducible P*_tet_* promoter. H_4_MPT, tetrahydromethanopterin; MFR, methanofuran; F_420_, coenzyme 420; Fd, ferredoxin; CoM-SH, coenzyme M; CoB-SH, coenzyme B; CoM-S-S-CoB, coenzyme M/coenzyme B heterodisulfide; CoA-SH, coenzyme A; CoFeSP, corrinoid iron-sulfur protein; AlsS, acetolactate synthase; AlsD, acetolactate dehydrogenase; pME2001, replicon for *M. thermautotrophicus*; *traJ*, gene encoding conjugal transfer function; ColE1, Gram-negative replicon for *Escherichia coli*; *catP*, chloramphenicol resistance gene; *neoR*, neomycin resistance gene; P*_hmtB_*, native histone promoter from *M. thermautotrophicus*; tMtaC, terminator from *M. acetivorans* (Pritchett and Metcalf, 2005); *alsD*, acetolactate decarboxylase gene from *S. thermophilus* codon-optimized for *M. thermautotrophicus*; *alsS*, acetolactate synthase gene from *S. thermophilus* codon-optimized for *M. thermautotrophicus*; P*_tet_*, anhydrotetracycline-inducible promoter; underlined, TATA motif; bold, *tetO* operator sequence; blue and bold, ribosomal binding site. Metabolic pathways native to *M. thermautotrophicus* are displayed in black, heterologous pathways are shown in orange, substrates hydrogen (H_2_) and carbon dioxide (CO_2_) are displayed in blue, the product methane (CH_4_) is shown in grey.

Our results provide a basis for enhancing acetoin bioproduction and for enabling the biosynthesis of additional value-added recombinant products from H₂ and CO₂, using *Methanothermobacter* species as robust and biotechnologically proven biocatalysts. This process can be aided by our genome-scale metabolic model (Casini et al., 2023) and enzyme engineering (*e.g.*, to overcome feedback inhibition of pathway enzymes), which recently was demonstrated in *M. marburgensis* (Unger et al., 2025).

## 2. Methods

### 2.1 Microbial strains and cultivation

All microbial strains and plasmids used in this study are listed in **Table 1**. Cloning of plasmids was performed in *Escherichia coli* NEB stable, whereas *E. coli* S17-1 was the donor strain for conjugational DNA transfer into *M. thermautotrophicus*. *E. coli* strains were cultivated in Luria-Bertani (LB) medium (Green and Sambrook, 2012) at 37°C and under constant shaking (150 rpm) for sufficient oxygenation. *M. thermautotrophicus* strains were cultivated anaerobically in mineral salt (MS) medium at 60°C when not otherwise stated or at 42°C (acetoin production) in serum bottles or bioreactors as previously described (Casini et al., 2023; Fink et al., 2021). Recombinant strains were selected by the addition of respective antibiotics to the following concentrations: 100 µg mL^-1^ carbenicillin, 30 µg mL^-1^ chloramphenicol, 250 µg mL^-1^ neomycin, and 10 µg mL^-1^ trimethoprim. Bottle experiments with recombinant *M. thermautotrophicus* strains were performed in 50 or 100 mL MS medium with neomycin in 250- or 1000-mL bottles, respectively, and under shaking conditions (150 rpm). Strains were provided with H_2_/CO_2_ as the sole energy and carbon source. For this purpose, the headspace of the bottles was filled with an H_2_/CO_2_ (80:20 vol-%) atmosphere to 1 bar overpressure. During the acetoin production experiments in batch cultivation, the headspace was refilled with H_2_/CO_2_ gas upon substrate depletion (monitored through headspace pressure) in 250-mL serum bottles and completely exchanged by five cycles of alternating vacuum application and gas purging to replace the produced CH_4_ in 1000-mL bottles, respectively. Because both β-galactosidase and acetoin genes were under the control of the P*_tet_* promoter, gene expression was induced using aTc after approximately 1.5 doublings.

**Table 1.**
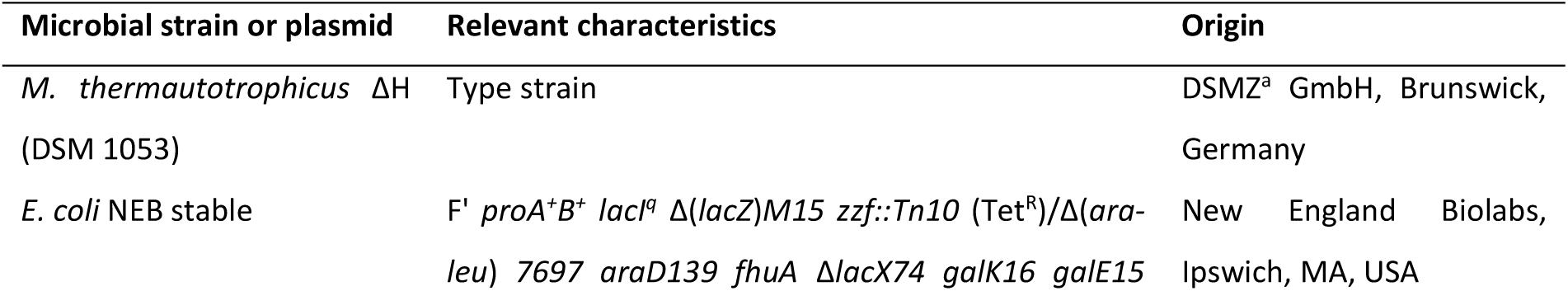
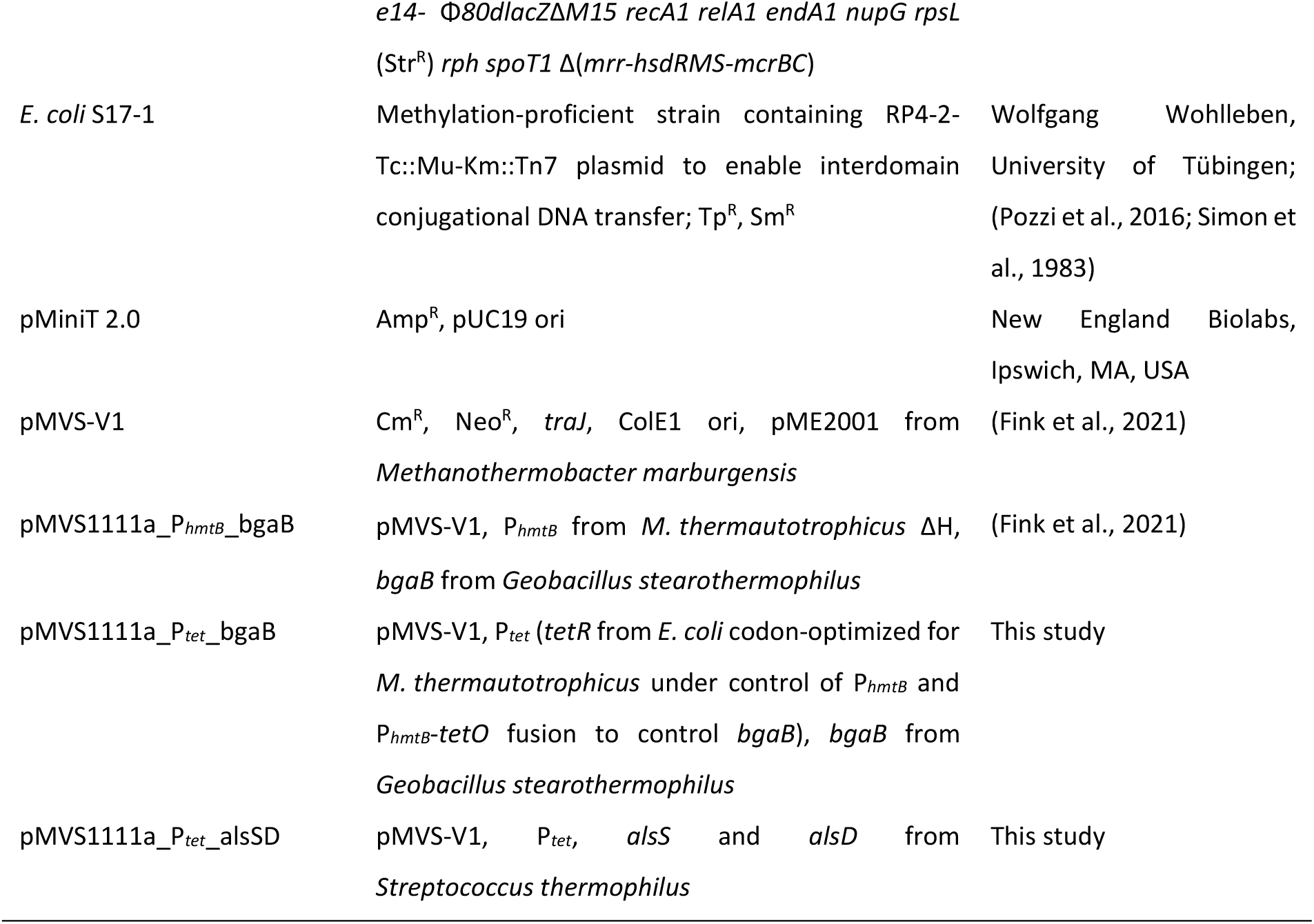
Microbial strains and plasmids used in this study.

### 2.2 Plasmid construction and DNA transfer

Standard molecular techniques were applied for plasmid construction. All primers and gBlock DNA fragments used in this study were synthesized by Integrated DNA Technologies (Coralville, IA, USA) and are listed in **Table 2**. DNA fragments for cloning were generated *via* PCR using the Q5 High-Fidelity DNA Polymerase (New England Biolabs, Ipswich, MA, USA) according to the manufactureŕs instructions and subsequently purified using the QIAquick® PCR Purification Kit (Qiagen GmbH, Hilden, Germany). Cloning was performed *via* Gibson assembly using the NEBuilder® HiFi DNA Assembly Master Mix (New England Biolabs, Ipswich, MA, USA). Chemically competent *E. coli* NEB stable cells were transformed with assembly mixtures according to previously described procedures (Fink et al., 2021). Successful plasmid constructions were verified *via* Sanger sequencing (GENEWIZ Germany GmbH, Leipzig, Germany).

**Table 2.**
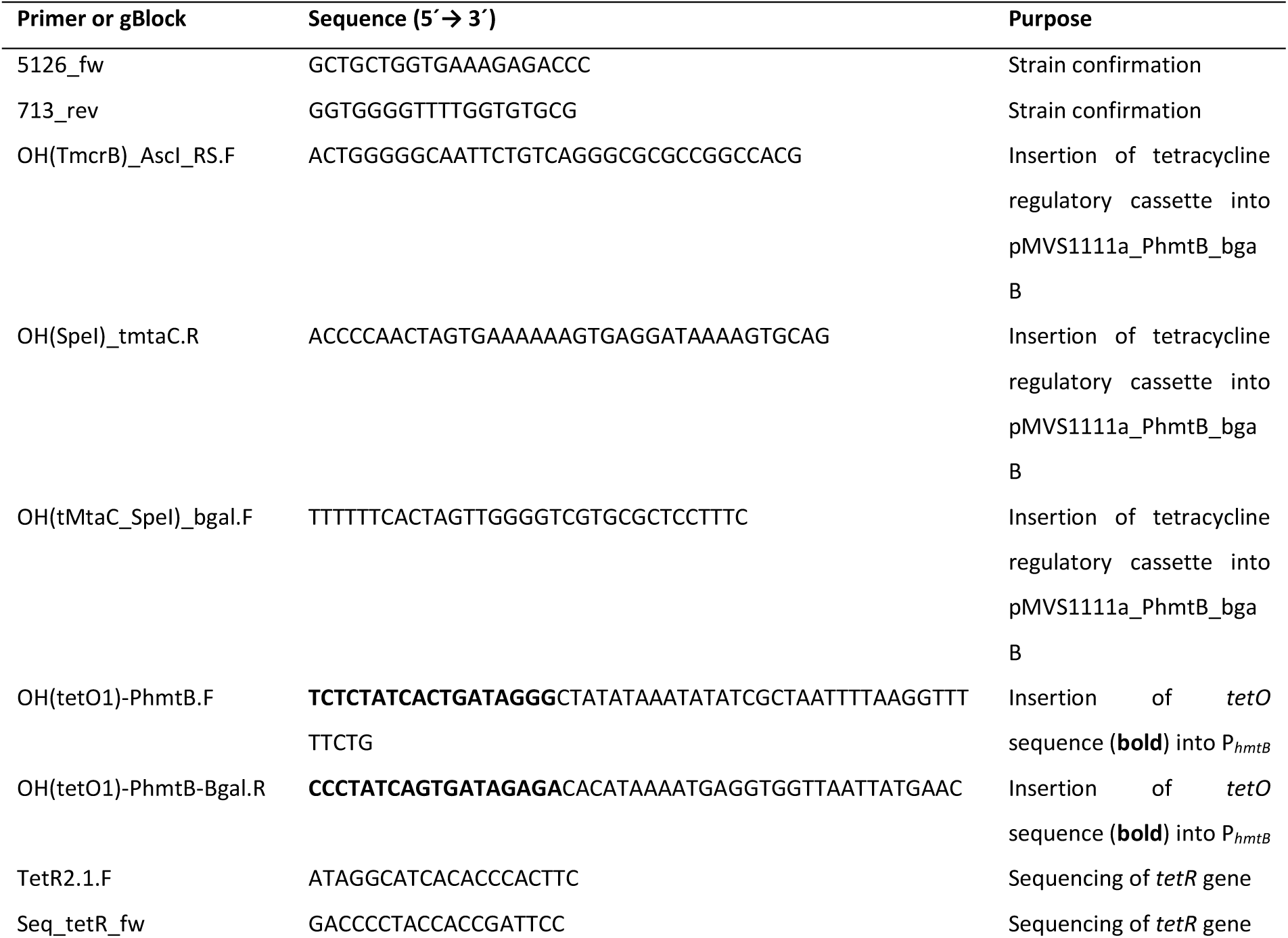
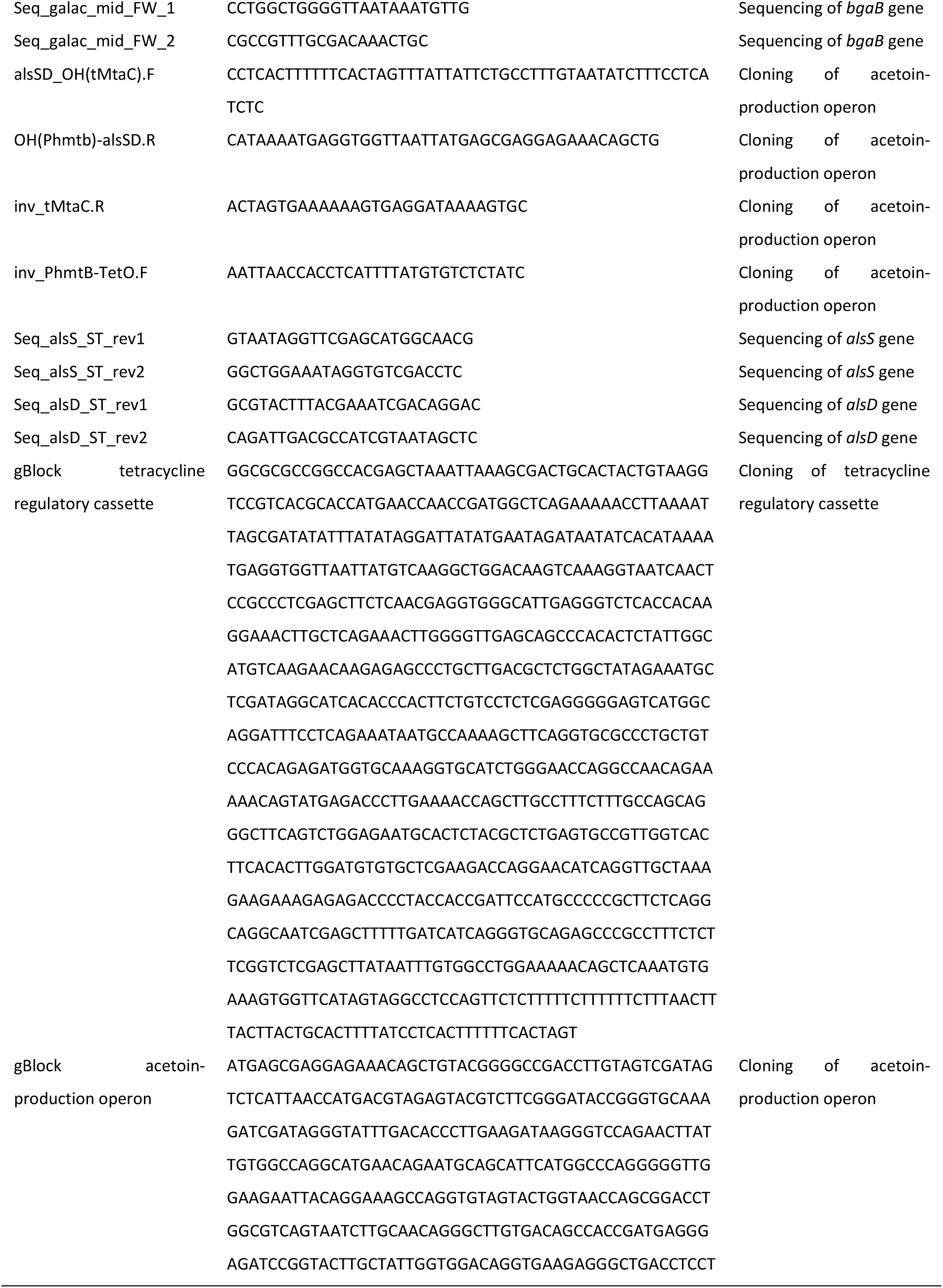
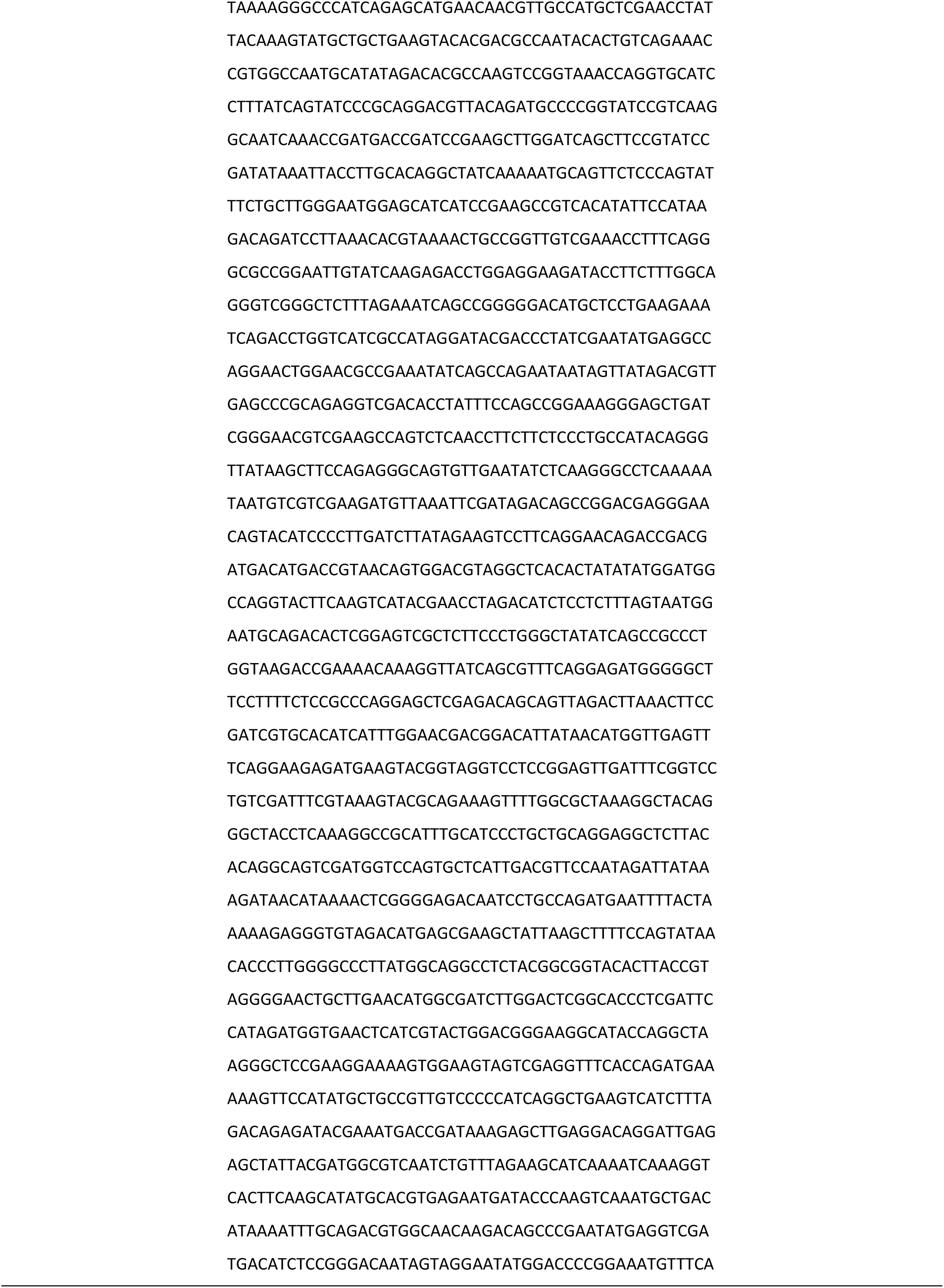
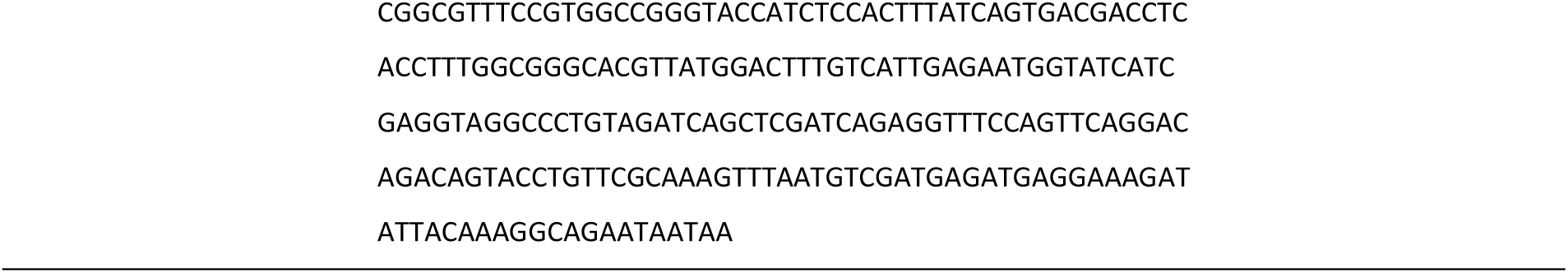
Primers and gBlock DNA fragments used in this study.

For the construction of plasmid pMVS1111a_P*_tet_*_bgaB, first, a tetracycline regulatory cassette composed of the transcriptional repressor gene sequence (*tetR*) from *E. coli* controlled by the P*_hmtB_* promoter and terminated by the tMtaC terminator from *Methanosarcina acetivorans* (Pritchett and Metcalf, 2005) was designed *in silico*. This tetracycline regulatory cassette was flanked by *Asc*I and *Spe*I restriction sites, codon-optimized for *M. thermautotrophicus*, and finally synthesized as a gBlock (Integrated DNA Technologies, Coralville, IA, USA; **Table 2**). The gBlock was subcloned into the pMiniT 2.0 vector and subsequently ligated into pMVS1111a_P*_hmtB_*_bgaB *via* Gibson assembly using the NEBuilder® HiFi DNA Assembly Master Mix (New England Biolabs, Ipswich, MA, USA). The primers OH(TmcrB)_AscI_RS.F and OH(SpeI)_tmtaC.R were used to amplify the tetracycline regulatory cassette, and OH(tMtaC_SpeI)_bgal.F and TmcrB.R were used to amplify pMVS1111a_P*_hmtB_*_bgaB backbone (**Table 2**). To put the *bgaB* gene under repression mediated by the TetR repressor, an operator sequence (*tetO1*; 5′-CCCTATCAGTGATAGAGA-3′) was inserted downstream of the TATA box of the P*_hmtB_* promoter controlling *bgaB* transcription following the design from Guss et al. (2008). This was achieved by amplification of pMVS1111a_P*_hmtB_*_bgaB containing the tetracycline regulatory cassette with primers OH(tetO1)-PhmtB.F and OH(tetO1)-PhmtB-Bgal.R (**Table 2**), and subsequent circularization of the amplified DNA fragment *via* created homologous overhangs using the NEBuilder® HiFi DNA Assembly Master Mix (New England Biolabs, Ipswich, MA, USA). The resulting plasmid was pMVS1111a_P*_tet_*_bgaB. To construct the acetoin-production plasmid, pMVS1111a_P*_tet_*_bgaB was used as the backbone vector, and the *bgaB* gene was exchanged against an acetoin-production operon, which consists of genes encoding an acetolactate synthase (*alsS*) and an acetolactate decarboxylase (*alsD*), both originating from *Streptococcus thermophilus*. The acetoin-production operon was codon-optimized for *M. thermautotrophicus*, synthesized as a gBlock, and subcloned into the pMiniT 2.0 vector. Subsequently, the acetoin-production operon was amplified from pMiniT 2.0 using primers alsSD_OH(tMtaC).F and OH(Phmtb)-alsSD.R, thereby, creating overhangs to the backbone vector pMVS1111a_P*_tet_*_bgaB, which was amplified with primers inv_tMtaC.R and inv_PhmtB-TetO.F to exclude the *bgaB* gene. Resulting DNA fragments were merged *via* Gibson assembly using the NEBuilder® HiFi DNA Assembly Master Mix (New England Biolabs, Ipswich, MA, USA), yielding the acetoin production plasmid pMVS1111a_P*_tet_*_alsSD.

All constructed plasmids were transferred into *E. coli* S17-1 and subsequently into *M. thermautotrophicus* according to the procedures described by Fink et al. (2021) with slight modifications. *M. thermautotrophicus* cells were cultivated overnight until early-stationary phase (OD_600_ approximately 0.3), transferred into 50-mL Falcon tubes inside the anaerobic chamber, and then harvested *via* centrifugation outside the anaerobic chamber at room temperature (10 min at 3,684 x g). Subsequently, cells were resuspended in 600 µL of the supernatant medium inside the anaerobic chamber. 300 µL of this concentrated *M. thermautotrophicus* suspension was carefully mixed with the pellet of 10 mL *E. coli* S17-1 harboring the plasmid to be transferred into *M. thermautotrophicus*. Afterward, 100 µL of the *M. thermautotrophicus*-*E. coli* S17-1 suspension was spotted on LB-MS agar plates for mating overnight at 37°C as otherwise previously described by Fink et al. (2021). LB-MS agar plates were prepared by adding peptone and yeast extract to MS medium during media preparation. Finally, recombinant strains were verified *via* PCR (Phire Plant Direct PCR Master Mix, Thermo Fisher Scientific, Waltham, MA, USA) using liquid cultures (Fink et al., 2021) and primers 5126_fw and 713_rev (**Table 2**).

### 2.3 β-galactosidase activity assays

For determining the optimal induction conditions of the P*_tet_* promoter, a thermostable β-galactosidase from *Geobacillus stearothermophilus* was used as a reporter protein (Fink et al., 2021). *M. thermautotrophicus* [pMVS1111a_P*_tet_*_bgaB] as well as *M. thermautotrophicus* [pMVS1111a_P*_hmtB_*_bgaB] (positive control) and *M. thermautotrophicus* [pMVS-V1] (negative control) were inoculated into 50 mL MS medium to an initial OD_600_ of 0.05 and cultivated at 60°C. After 1.5 doubling times, *M. thermautotrophicus* [pMVS1111a_P*_tet_*_bgaB] cultures were induced with varying concentrations of aTc, while one culture remained uninduced to determine the basal activity of the P*_tet_* promoter. To determine the β-galactosidase enzyme activity, 4 mL of culture were withdrawn 14, 38, 63, and 135 h after induction and centrifuged (10 min, 3,684 x g, 4°C). Subsequently, cells were washed twice with ice-cold phosphate-buffered saline (137 mM NaCl, 10 mM Na_2_HPO_4_ x 2 H_2_O, 2 mM KH_2_PO_4_, 2.7 mM KCl, pH 7.4) before suspending cell pellets in 200 µL cold lysis buffer (100 mM Tris, 20 % (v/v) glycerol, 1 mM dithiothreitol (DTT), pH 8, 1 mM phenylmethylsulfonyl fluoride [PMSF, added freshly before use from 100 mM stock solution in isopropanol]). For cell lysis, the suspension was filled into 2-mL cryotubes with approximately 200 µL glass beads (0.1 mm diameter) and then treated in the FastPrep-24^TM^ 5G bead beater (MP Biomedicals, Santa Ana, CA, USA) for 5 cycles à 40 s and 6 m/s with intermittent cooling on ice for 1 min between cycles. Cell-free extracts were harvested by centrifugation at 10,000 x g for 10 min and 4°C. 10 µL of cell extract was added to 60 µL substrate solution (60 mM Na_2_HPO_4_ x 2 H_2_O, 40 mM NaH_2_PO_4_ x H_2_O, 10 mM KCl, 50 mM β-mercaptoethanol, 3.3 mM o-nitrophenyl-β-D-galactopyranoside [ONPG] (Jensen et al., 2017)), supplied in a 96-well microtiter plate (Cole-Parmer, Vernon Hills, IL, USA). The microtiter plate was incubated at 60°C for 2 h before the β-galactosidase reaction was terminated by the addition of 70 µL stop solution (1 M Na_2_CO_3_). Afterwards, the absorption at 420 nm (A_420_) was measured using the Victor Nivo^TM^ Multimode plate reader (PerkinElmer, Waltham, MA, USA). For the determination of the specific β-galactosidase activity, the protein concentration in the cell extracts was measured using the Qubit^TM^ Protein Assay Kit in the Qubit^TM^ Flex Fluorometer (Thermo Fisher Scientific, Waltham, MA, USA) following manufactureŕs instructions. Finally, specific β-galactosidase activities (*a* in nmol min^-1^ mg protein^-1^) were calculated with the following equation 1 (Sedlak et al., 1995):

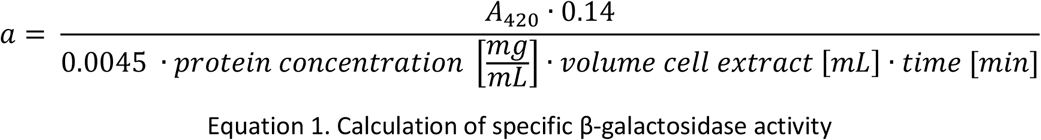

with factor 0.14 correcting for the reaction volume, 0.0045 being the A_420_ of a 1 nmol mL^-1^ solution of *o*-nitrophenol (product of the β-galactosidase reaction), a cell extract volume of 0.01 mL, and a reaction time of 120 min.

### 2.4 Bioreactor experiments

Stainless-steel bioreactor systems (Sartorius Stedim Biotech GmbH, Göttingen, Germany) with a working volume of 2 L were used for the cultivation of recombinant *M. thermautotrophicus*. The medium for bioreactor studies was prepared as previously described by Casini et al. (2023) and additionally contained 250 µg mL^-1^ neomycin. The medium was prepared as a 10x stock solution, diluted with demineralized water prior to use, and autoclaved directly in the bioreactor vessel, as detailed below. Media feed and Na_2_S supplementation, pH, gas sparging, and temperature were controlled using the bioreactor control tower and software (Biostat B-DCU and BioPAT, Sartorius Stedim Biotech GmbH, Göttingen, Germany). The pH was maintained at 7.3 with 5 M NaOH and 2 M H_2_SO_4_ solutions. An 80:20 vol-% H_2_/CO_2_ gas mixture was supplied using separate mass flow controllers (Alicat MC Series, Alicat Scientific, Tucson, AZ, USA) at 25 or 50 mL min^-1^. The H_2_/CO_2_ gas was supplied directly into the bioreactor, and an additional N_2_ flow (100 mL min^-1)^ -gas in a mixing chamber (approximately 100 mL) to create a combined flow to the off-gas mass spectrometer (PrimaBT Benchtop Process Mass Spectrometer, Thermo Fisher Scientific, Waltham, MA, USA) for determination of N_2_, H_2_, CO_2_, and CH_4_ concentrations. To correct for fluctuations in the off-gas measurements due to a small backpressure in the mass spectrometer, a five-point moving average was calculated from the off-gas data before gas productivities were calculated.

#### 2.4.1 Fed-batch operation

Before starting the experiment, the bioreactors, containing 2 L of cultivation medium, were autoclaved at 121°C and 1.2 bar pressure for 20 min. After autoclaving, the bioreactors were sparged with N_2_ overnight at 60°C to ensure anaerobic conditions. Next, final concentrations of 250 µg mL^-1^ neomycin for plasmid selection, as well as 0.5 g L^-1^ anaerobic L-cysteine-HCl and 0.3 g L^-1^ Na_2_S x 9 H_2_O to reduce the medium were added, stirring was set to 100 rpm, and the pH control was started. Once the bioreactor was reduced (redox potential at least -300 mV), the inlet gas was switched to a continuous supply of 25 mL min^-1^ H_2_/CO_2_, the N_2_ was connected to the gas mixing chamber with the off-gas, and the bioreactor was inoculated with 20 mL of an actively growing pre-culture of *M. thermautotrophicus* [pMVS-V1] or *M. thermautotrophicus* [pMVS1111a_P*_tet_*_alsSD], respectively. After an overnight growth phase at 60°C, the temperature was decreased to 42°C and the stirring speed increased to 250 rpm. When an OD_600_ of 0.1 was reached, acetoin production was induced with 1 µg mL^-1^ aTc, and the stirring speed was increased to 500 rpm. After 8 h at 500 rpm, the stirring speed was further increased to 750 rpm. To assess whether the acetoin production could be further boosted, the stirring speed was increased to 1000 rpm after 75 h of operation, and the H_2_/CO_2_ supply was increased to 50 mL min^-1^ after 92 h of operation. The fed-batch experiments were terminated after a 10-day operating period.

#### 2.4.2 Chemostat operation

To assess the potential for continuous acetoin production, a chemostat bioreactor experiment with the acetoin producer *M. thermautotrophicus* [pMVS1111a_P*_tet_*_alsSD] was carried out. The chemostat was started as described for fed-batch; initial batch growth occurred at 60°C. During the batch phase, the H_2_/CO_2_ supply was cut off twice due to technical issues, leading to a longer 60°C phase of 2 days instead of overnight. During batch operation, the stirring speed was ramped up gradually from 100 to 500 rpm. After reaching an OD of 0.1, the temperature was reduced to 42°C and the bioreactor was switched to continuous mode with a hydraulic retention time (HRT) of 1 day, amounting to a medium inflow rate of 83.3 mL h^-1^. Na_2_S x 9 H_2_O was continuously added from a 111 g L^-1^ stock solution at a rate of 0.255 mL h^-1^ to maintain a reduced environment and provide sufficient sulfur for growth. A stable biomass concentration was reached after 3 days in continuous mode after which acetoin production was induced with 1 µg mL^-1^ aTc. The acetoin production genes originate from *S. thermophilus*, which is a microbe with an optimum growth temperature of around 35-42°C (Radke-Mitchell and Sandine, 1986). However, the production host *M. thermautotrophicus* has an optimum growth temperature of 60°C to 65°C. To assess the influence of temperature on acetoin production, the chemostat culture was exposed to a temperature gradient. After induction, the bioreactor was maintained at 42°C for 3 HRTs. Then, technical disturbances were experienced for 3 HRTs. After stabilizing the bioreactor for 1 HRT, the temperature was increased to 50°C for 4 HRTs, and subsequently to 60°C for 3 HRTs.

### 2.5 Analytical methods

Growth of *M. thermautotrophicus* strains was routinely monitored by measuring the optical density at 600 nm (OD_600_) using the BioMate^TM^ 160 UV-Vis spectrophotometer (Thermo Fisher Scientific, Waltham, MA, USA). For the analysis of the acetoin content, 2-mL culture samples were withdrawn from *M. thermautotrophicus* cultures in regular intervals and centrifuged (10 min, 3,684 x g, 4°C). The supernatant was filtered using 0.22 µm PVDF syringe filters (13 mm, Carl Roth GmbH & Co. KG, Karlsruhe, Germany) and then analyzed *via* gas chromatography (GC). For that purpose, 500 µL of filtered supernatant were filled into 2-mL vials (VWR International GmbH, Darmstadt, Germany) and closed with screw caps. Prepared samples were analyzed using the 7890B GC system (Agilent Technologies, Santa Clara, CA, USA) equipped with a DB-FATWAX UI capillary column (inner diameter 0.25 mm x 30 m; 0.25 µm film) and a flame ionization detector set to 250°C. H_2_ served as the carrier gas with a column flow of 21.4 mL min^-1^. The injection temperature was set to 200°C, the detector gases were H_2_ (35 mL min^-1^) and air (350 ml min^-1^). 0.5 µL of prepared sample were injected into the GC and analyzed using the following oven program: 80°C for 0.5 min, followed by a gradual temperature increase to 180°C at 20°C min^-1^, finally 180°C were held for 1 min. External standards with defined acetoin concentrations were analyzed for calibration purposes.

To confirm the identity of the produced compound as acetoin, gas chromatography coupled with mass spectrometry (GC-MS) was carried out after bottle experiments in 100 mL MS medium. Culture samples were centrifuged and filtered as described for GC analysis and then underwent a liquid-liquid extraction procedure to generate water-free samples for the MS analysis. For that purpose, 250 µL ethyl acetate were added to 1 mL of filtered supernatant for the extraction of acetoin. Subsequently, samples were vortexed for 3 x 3 min, then incubated for 20 min at room temperature under constant shaking (1800 rpm), and finally centrifuged (10,000 x g, 2 min). The upper phase was transferred into 0.25-mL micro-inserts (Carl Roth GmbH & Co. KG, Karlsruhe, Germany) and then analyzed via GC-MS using the 7890B GC system (Agilent Technologies, Santa Clara, CA, USA) equipped with a DB-FATWAX UI capillary column (inner diameter 0.25 mm x 30 m; 0.25 µm film) and a 5977B mass selective detector (source temperature 230°C, quadrupole temperature 150°C). The carrier gas flow (H_2_) was 1.4 mL min^-1^, the injection temperature was set to 250°C. 1.8 µL of the processed water-free supernatant were injected into the GC-MS system and analyzed using the following temperature profile: 50°C for 3 min followed by a temperature increase to 85°C at 5°C min^-1^, a plateau at 85°C for 4 min followed by another temperature increase to 250°C at a rate of 35°C min^-1^, and a final plateau at 250°C for 4 min. External, defined acetoin standards were processed equally to the culture samples and used for peak identification. An MS scan was performed for acetoin peaks between 40 and 89 m/z to record the mass-to-charge ratio of generated ions of the identified peak. The obtained mass spectra were compared with the NIST database (National Institute of Standards and Technology, Gaithersburg, MD, USA) to identify acetoin.

### 2.6 Statistical Analyses

All batch experiments for the β-galactosidase activity assay and acetoin production were performed in biological triplicates (n=3). For the statistical analysis, no samples were removed from the dataset. The average volumetric acetoin-production rates in closed-batch and fed-batch fermentations were calculated as described in equation 2. Average specific acetoin production rates normalized to OD_600_ of closed-batch and fed-batch fermentation were calculated as in equation 3.

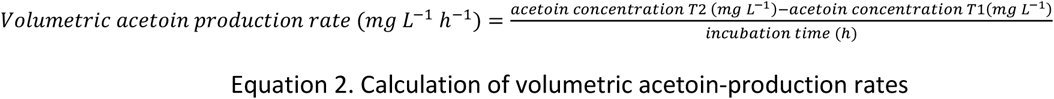

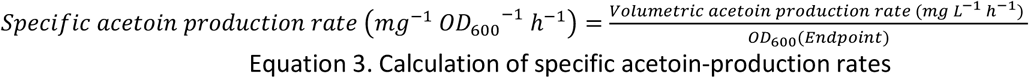

Before conducting statistical tests, the samples were analyzed for equal distribution with Shapiro-Wilk′s test and equal variance between data series with Levenés test. For the β-galactosidase activity assay, equal distribution and equal variance were given, therefore, a one-way analysis of variance (ANOVA) was performed to identify significant differences between the datasets. Subsequently, a Tukey honest significant difference test (HSD) was added to identify significantly differing data series.

In the 250-mL bottle batch samples for acetoin production, equal distribution but no equal variance was observed. Therefore, a Welch′s ANOVA was performed, which is not dependent on equal variance. Subsequently, a Tukey HSD was added to identify significantly differing data series. For the 1000-mL bottle batch samples for acetoin production, data sets with only two data series were collected. Here, statistical analysis with homoscedastic, one-sided Student′s t-test was performed. All tests were rendered significant when the p-value was below 0.05 and highly significant when the p-value was below 0.01.

## 3. Results

### 3.1 An anhydrotetracycline-inducible promoter system is functional in *M. thermautotrophicus*

Controlling gene expression is an important component of a genetic toolbox for metabolic engineering. We aimed to implement an inducible promoter, based on the *tetR*-P*_tet_* system, which is known to be active in *M. acetivorans* (Guss et al., 2008). It has been shown that the *tetR*-P*_tet_* system can be used for temperature-dependent induction in thermophilic microbes, because the TetR-binding is temperature-sensitive (Mol, 2022). This leads to the detachment of TetR from the DNA when the temperature is increased above approximately 60 °C, which was recently demonstrated for *M. marburgensis* as well (Unger et al., 2025). However, we used the AlsS and AlsD enzymes from *S. thermophilus*, which has a growth optimum of 42°C (Radke-Mitchell and Sandine, 1986), and to our knowledge functionality of these enzymes above 42°C was not tested yet (Akyol et al., 2015). Thus, here, we developed a system that can be induced by aTc at temperatures below 60°C. We constructed the strain *M. thermautotrophicus* [pMVS1111a_P*_tet_*_bgaB], which carries a replicative plasmid encoding a thermophilic β-galactosidase, and which is under the control of the *tetR*-P*_tet_* system (**Figure 1C**). *M. thermautotrophicus* [pMVS-V1] and *M. thermautotrophicus* [pMVS1111a_P*_hmtB_*_bgaB], which we constructed in previous work (Fink et al., 2021), were used as empty-vector and constitutive-expressing control strains.

We induced the expression of the *bgaB* gene with varying concentrations of aTc and measured the β-galactosidase activity (**Figure 2**). The growth (OD_600_) of the strains was not impacted by the tested aTc concentrations (**Figure 2A**). We observed a basal specific β-galactosidase activity of 9.74 ± 5.06 nmol min^-1^ mg_protein_^-1^ in the empty-vector control strain (*M. thermautotrophicus* [pMVS-V1]). In the uninduced *M. thermautotrophicus* [pMVS1111a_P*_tet_*_bgaB] strain, the activity increased to a maximum of 15.82 ± 3.79 nmol min^-1^ mg ^-1^ over time, compared to the basal activity of the empty-vector control strain. The constitutive-expressing *M. thermautotrophicus* [pMVS1111a_P*_hmtB_*_bgaB] strain showed a maximum activity of 45.58 ± 3.91 nmol min^-1^ mg_protein_^-1^. Induction with aTc in *M. thermautotrophicus* [pMVS1111a_P*_tet_*_bgaB] led to activities, which were similar to the constitutive-expressing strain shortly after aTc addition (**Figure 2**). While we observed induction with all tested aTc concentrations, the addition of 1 µg mL^-1^ resulted in the highest measured activity of 58.02 ± 3.17 nmol min^-1^ mg ^-1^, which was significantly higher with a 3.7-fold increase in activity compared to the uninduced control (**Figure 2C**). Induction with 1 µg mL^-1^ of aTc also resulted in a significantly higher specific β-galactosidase activity compared to all other aTc concentrations for induction and compared to the constitutively expressed positive control (**Figure 2C**). Specific β-galactosidase activity stayed relatively stable over the duration of the experiment, with slight variations (**Figure 2B**). Overall, our results demonstrate the successful implementation of an aTc-inducible promoter system in *M. thermautotrophicus* for controlling gene expression.

**Figure 2.**
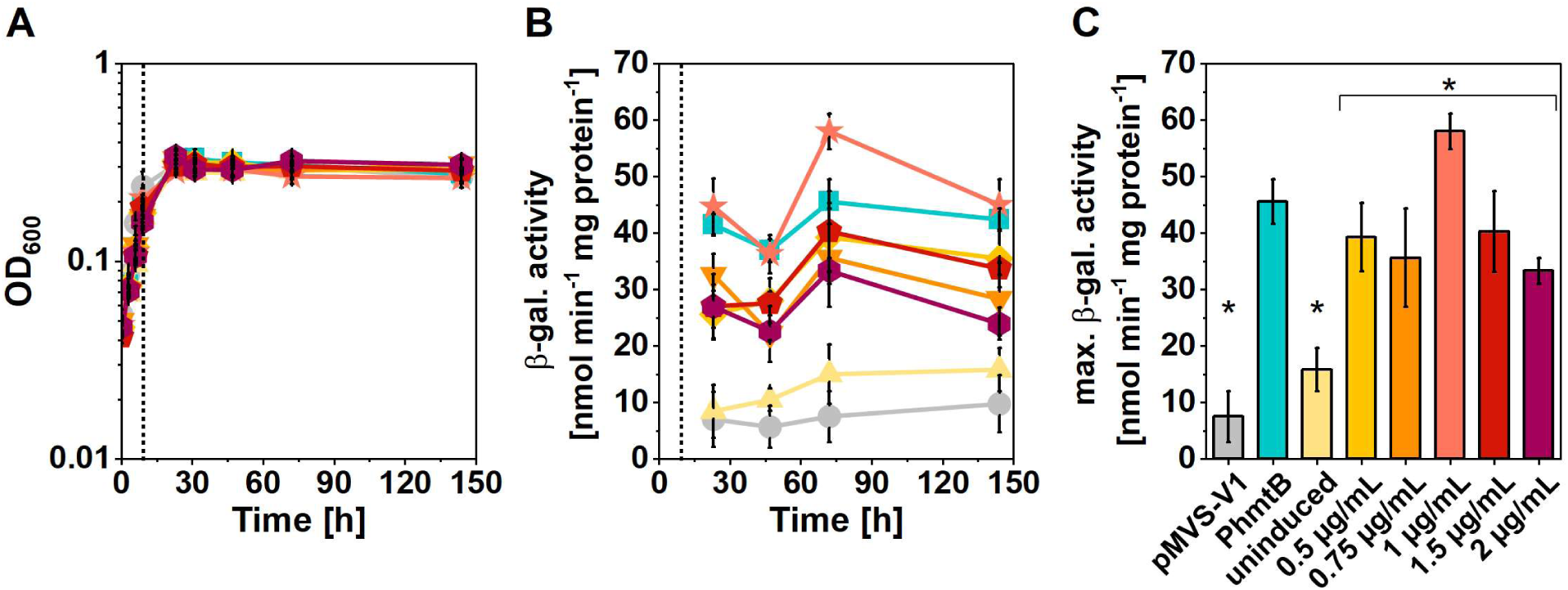
Characterization of P*_tet_* promoter in *M. thermautotrophicus* using a thermostable β-galactosidase (BgaB) as a reporter protein. A, Growth (OD_600_); **B**, β-galactosidase activity over time; **C**, maximum β-galactosidase activity. Expression of *bgaB* was induced using anhydrotetracycline (aTc; dotted line). Circles, *M. thermautotrophicus* [pMVS-V1]; squares, *M. thermautotrophicus* [pMVS1111a_P*_hmtB_*_bgaB]; triangles, *M. thermautotrophicus* [pMVS1111a_P*_tet_*_bgaB] uninduced; diamonds, *M. thermautotrophicus* [pMVS1111a_P*_tet_*_bgaB] + 0.5 µg mL^-1^ aTc; downward-facing triangles, *M. thermautotrophicus* [pMVS1111a_P*_tet_*_bgaB] + 0.75 µg mL^-1^ aTc; stars, *M. thermautotrophicus* [pMVS1111a_P*_tet_*_bgaB] + 1 µg mL^-1^ aTc; pentagons, *M. thermautotrophicus* [pMVS1111a_P*_tet_*_bgaB] + 1.5 µg mL^-1^ aTc; hexagons, *M. thermautotrophicus* [pMVS1111a_P*_tet_*_bgaB] + 2 µg mL^-1^ aTc. Error bars indicate standard deviations, n=3. The single asterisks indicate significant difference from the group below the bracket with asterisk (p<0.05, α=0.05). 1 µg mL^-1^ is significantly different from the other induced samples, which is not indicated by an additional symbol.

### 3.2 Recombinant acetoin production can be achieved in *M. thermautotrophicus*

Next, we aimed at acetoin production with a plasmid-encoded acetoin-production operon, consisting of the *alsS* and *alsD* genes from *S. thermophilus*, which were codon-optimized for *M. thermautotrophicus*, and controlled by the aTc-inducible promoter system (**Figure 1**). We compared growth and acetoin production of the strain *M. thermautotrophicus* [pMVS1111a_P*_tet_*_alsSD] carrying the acetoin-production operon with the wildtype and empty plasmid-carrying strain. We used 42°C as the incubation temperature, because this allows good growth of *M. thermautotrophicus* while displaying the optimum growth temperature of *S. thermophilus*, which should increase the likelihood of functional enzyme production from the acetoin-production operon. We speculated that during growth, flux redistribution into acetoin production may not occur, because of regulation of biomass biosynthesis to avoid loss of energy and carbon. Thus, we initially performed batch experiments with gas refeeding. In batch experiments in 250-mL serum bottles, all strains grew comparably, while growth was not steady due to the fast depletion of substrate gases that we regularly refilled (**Figure 3A-C**). However, after induction with aTc, *M. thermautotrophicus* [pMVS1111a_P*_tet_*_alsSD] achieved a final acetoin titer of 0.117 ± 0.007 mM and a volumetric production rate of 0.540 ± 0.047 µmol L^-1^ h^-1^ (**Figure 3C**, **Table 3**). Surprisingly, we observed the slow accumulation of acetoin until the end of the experiment also in all controls, however, at ∼10-fold and significantly lower final titers for the controls (**Figure 3C**, **Table 3**). In this experiment, the limited headspace-to-liquid ratio (5:1) and sampling artifacts due to withdrawal of relatively high liquid volumes compared to total volumes prompted us to perform another batch experiment in 1000-mL bottles with a headspace-to-liquid-ratio of 10:1, while omitting the wildtype control (**Figure 3D-F**, **Table 3**). In this second batch experiment, we fully replaced the headspace after each depletion of the substrate gas, to replace the produced CH_4_. All strains grew comparably with a clear phase of increasing OD_600_ followed by a plateau (slight decrease in OD_600_), during which the substrate gases were still consumed efficiently (**Figure 3D,E**). In the first 68 h after induction, the production of acetoin for *M. thermautotrophicus* [pMVS1111a_P*_tet_*_alsSD] was aTc-dependent, and we did not detect acetoin in the controls (**Figure 3**). However, after growth ceased in all cultures and stagnation of the OD_600_ occurred, the acetoin concentrations in all cultures, including the controls, continued to increase, but at a lower rate that was similar for all cultures, independent of the presence of the acetoin-production operon and induction (**Figure 3**, **Table 3**). During this phase, we did not detect statistical differences in the volumetric acetoin-production rates among all tested strains. This non-specific increase of acetoin was likely due to chemical decomposition of acetolactate, which accumulated under non-growth conditions with sufficient gas supply, as discussed later. 68 h after induction, the induced *M. thermautotrophicus* [pMVS1111a_P*_tet_*_alsSD] had produced 0.259 ± 0.033 mM acetoin (Phase 1, 3.814 ± 0.478 µmol L^-1^ h^-1^). Eventually, this strain reached a final titer of 0.447 ± 0.082 mM acetoin (Phase 2, 0.780 ± 0.216 µmol L^-1^ h^-1^), which was a ∼2.4-fold higher final titer than in the controls. Compared to the first experiment, the final acetoin titer increased by 380% for the induced *M. thermautotrophicus* [pMVS1111a_P*_tet_*_alsSD] strain. From this experiment, we confirmed acetoin by GC-MS (**Supplemental Figure 1**).

**Figure 3.**
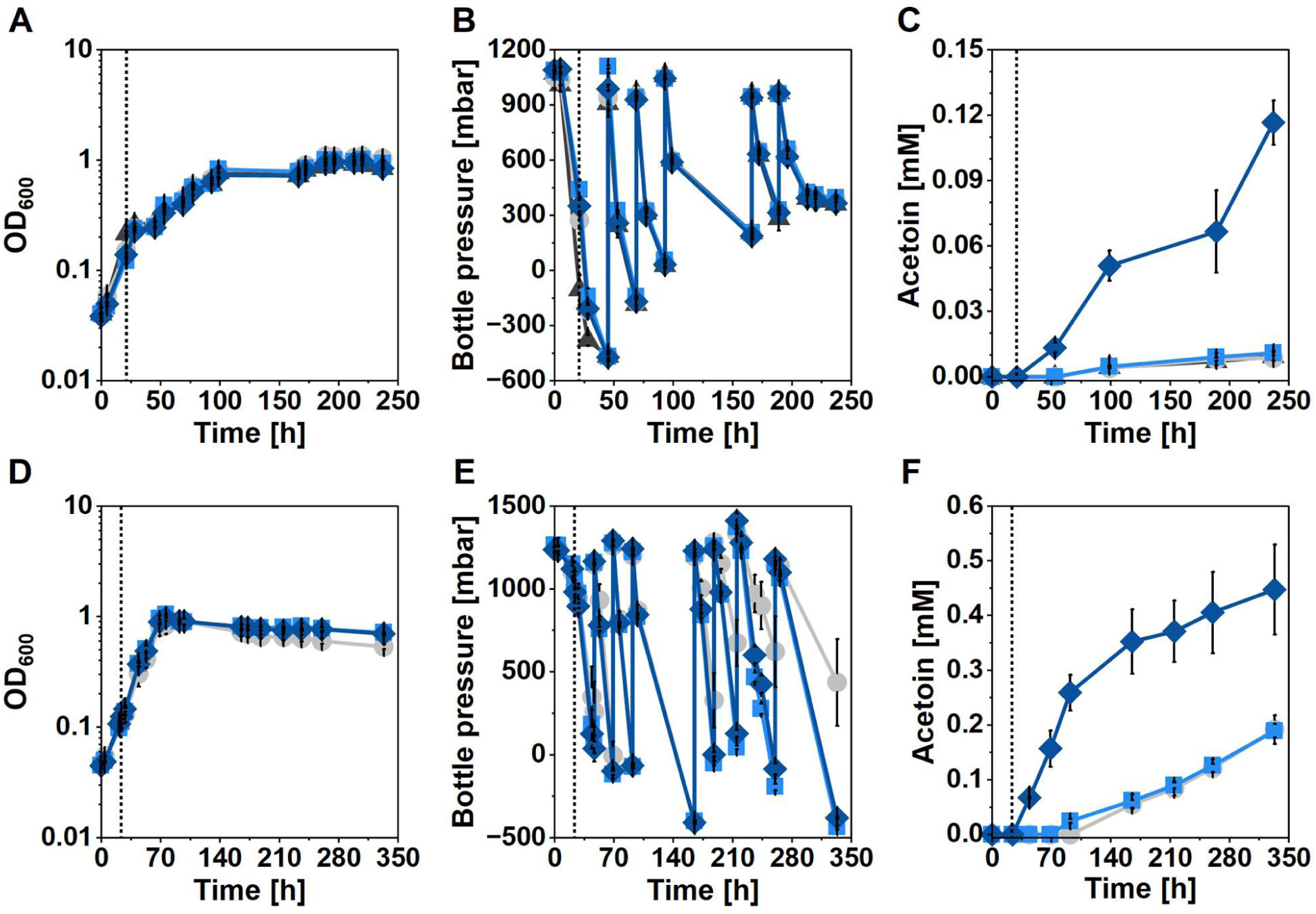
**Recombinant acetoin production with *M. thermautotrophicus*. A-C**, Cultivation in 50 mL MS medium (250-mL serum bottles); **D-F**, Cultivation in 100 mL MS medium (1000-mL bottles). Black triangles, *M. thermautotrophicus* wildtype; grey circles, *M. thermautotrophicus* [pMVS-V1]; light blue squares, *M. thermautotrophicus* [pMVS1111a_P*_tet_*_alsSD] uninduced; dark blue diamonds, *M. thermautotrophicus* [pMVS1111a_P*_tet_*_alsSD] induced using 1 µg mL^-1^ anhydrotetracycline (dotted line). Left panels (**A** and **D**), growth (OD_600_); middle panels (**B** and **E**), bottle pressure in headspace; right panels (**C** and **F**), acetoin concentration. All experiments were carried out at 42°C. The headspace of the bottles was refilled with fresh H_2_/CO_2_ upon depletion (**A-C**) or completely exchanged (**D-E**). Error bars indicate standard deviations, n=3.

**Table 3.** Summary of growth and acetoin productivity across the different bottle experiments.

| Strain | final OD<sub>600</sub> | Acetoin [mM] | Production rate<br>[μmol L<sup>-1</sup> h<sup>-1</sup>] | Acetoin<br>[mg L<sup>-1</sup>] | Production rate<br>[mg L<sup>-1</sup> h<sup>-1</sup>] |
| --- | --- | --- | --- | --- | --- |
| **250-mL serum bottles** |  |  |  |  |  |
| wildtype | 0.84 ± 0.02 | 0.009 ± 0.002 | 0.043 ± 0.010 | 0.822 ± 0.183 | 0.004 ± 0.001 |
| pMVS-V1 | 1.04 ± 0.04 | 0.009 ± 0.001 | 0.040 ± 0.006 | 0.758 ± 0.107 | 0.004 ± 0.001 |
| uninduced | 0.94 ± 0.03 | 0.011 ± 0.001 | 0.050 ± 0.005 | 0.960 ± 0.102 | 0.004 ± 0.001 |
| induced | 0.84 ± 0.02 | 0.117 ± 0.010 | 0.540 ± 0.047 | 10.279 ± 0.900 | 0.047 ± 0.004 <sup>1</sup> |
| **1000-mL bottles** |  |  |  |  |  |
| **Phase 1 (24-92 h)** |  |  |  |  |  |
| pMVS-V1 | 0.80 ± 0.13 | 0.000 ± 0.000 | 0.000 ± 0.000 | 0.000 ± 0.000 | 0.000 ± 0.000 <sup>2</sup> |
| uninduced | 0.95 ± 0.08 | 0.025 ± 0.004 | 0.367 ± 0.062 | 2.197 ± 0.374 | 0.032 ± 0.005 |
| induced | 0.89 ± 0.05 | 0.259 ± 0.033 | 3.814 ± 0.478 | 22.848 ± 2.865 | 0.336 ± 0.042 <sup>1</sup> |
| **Phase 2 (92-333 h)** |  |  |  |  |  |
| pMVS-V1 | 0.53 ± 0.03 | 0.192 ± 0.026 | 0.795 ± 0.108 | 16.887 ± 2.288 | 0.070 ± 0.010 <sup>3</sup> |
| uninduced | 0.71 ± 0.01 | 0.189 ± 0.012 | 0.682 ± 0.056 | 16.681 ± 1.087 | 0.060 ± 0.005 <sup>3</sup> |
| induced | 0.70 ± 0.02 | 0.447 ± 0.082 | 0.780 ± 0.216 | 39.412 ± 7.225 | 0.069 ± 0.019 <sup>3</sup> |

<sup>1</sup> Significant difference to all other sample series in the section
<sup>2</sup> No statistical analysis performed due to absence of product
<sup>3</sup> No significant differences among samples of Phase 2 (p>0.05)

### 3.3 Acetoin production is facilitated in fed-batch and chemostat bioreactors

Based on the observation that recombinant acetoin production was growth-coupled and non-specific acetoin accumulation was found when growth ceased, we argued that in a batch experiment with continuous gas feeding (fed-batch), and thus more sustained growth, we would be able to increase the acetoin production further. Therefore, we performed a fed-batch bioreactor experiment with a single bioreactor for the *M. thermautotrophicus* [pMVS-V1] and two replicates of the *M. thermautotrophicus* [pMVS1111a_P*_tet_*_alsSD] strain (**Figure 4, Supplemental Figure 2,** **Table 4**). Indeed, we observed growth of all cultures throughout the experiment, which slowed down from 150 h but did not cease (**Supplemental Figure 2**). Accordingly, we did not measure any acetoin for the control strain at any point of the experiment. Surprisingly, one replicate of the *M. thermautotrophicus* [pMVS1111a_P*_tet_*_alsSD] strain reached a much higher final OD_600_ of 1.72 compared to the second replicate for this strain (0.93) and the control strain (0.95), which both showed a similar growth and CH_4_ production profile. The higher OD_600_ resulted in higher CH_4_ and acetoin production (**Supplemental Figure 2**) but this was comparable to both other bioreactors (control and production strain) in terms of CH_4_ production and to the second replicate of the production strain in terms of acetoin production, when normalized to the OD_600_ (**Figure 4**). The final average specific acetoin titer (normalized to OD_600_) in the fed-batch bioreactors was 0.367 ± 0.338 mmol L^-1^ OD_600_^-1^ (**Table 4**), which is slightly higher compared to the 1000-mL bottle experiment for Phase 1 (**Table 3**). The achieved final acetoin titer and specific production rate did not increase compared to the overall 1000-mL bottle experiment (**Tables 3, 4, and 5**). However, those values were reached after a 250-h operation period in the fed-batch bioreactors compared to 333 h in the 1000-mL bottle experiment. Further, we did confirm that non-specific acetoin accumulation in the control strain did not occur during growth for fed-batch conditions.

**Figure 4.**
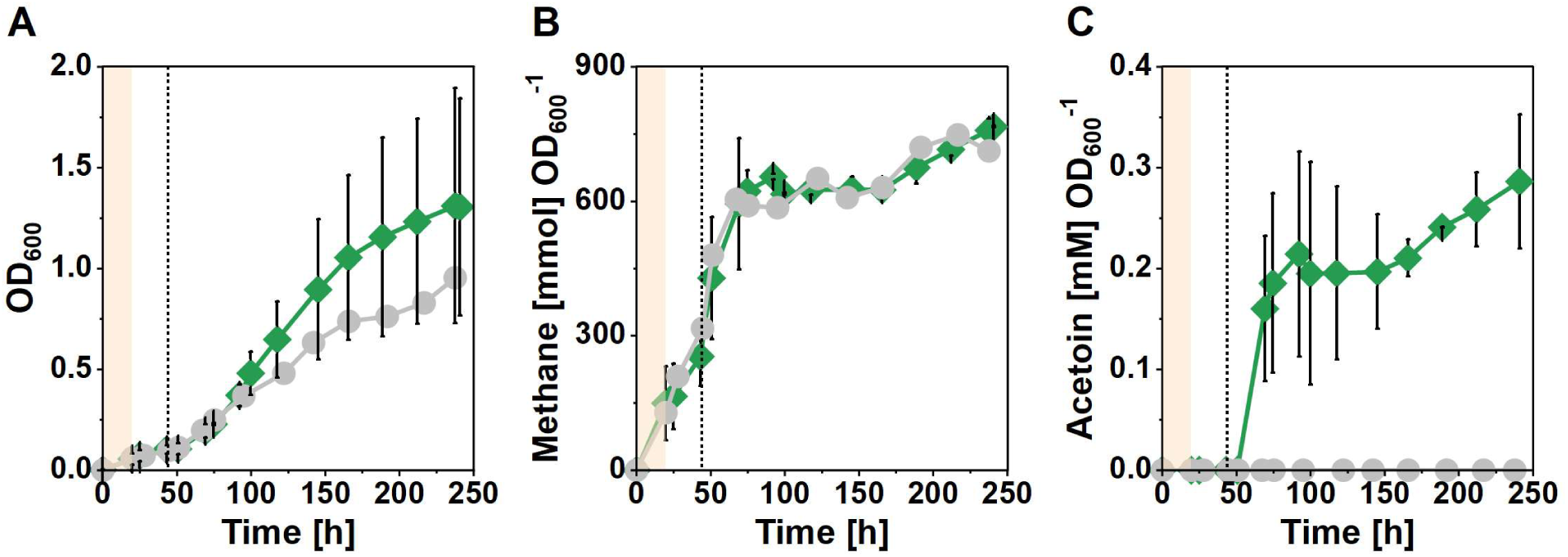
Recombinant acetoin production with *M. thermautotrophicus* in fed-batch bioreactors. **A**, Growth, OD_600_; **B**, normalized cumulative methane production; **C**, normalized acetoin production. Grey circles, *M. thermautotrophicus* [pMVS-V1] (n=1); green diamonds, *M. thermautotrophicus* [pMVS1111a_P*_tet_*_alsSD] (n=2). Initial growth occurred at 60°C (light orange shading), afterwards, the temperature was set to 42°C, and acetoin production was induced with 1 µg mL^-1^ anhydrotetracycline (dotted line).

**Table 4.**
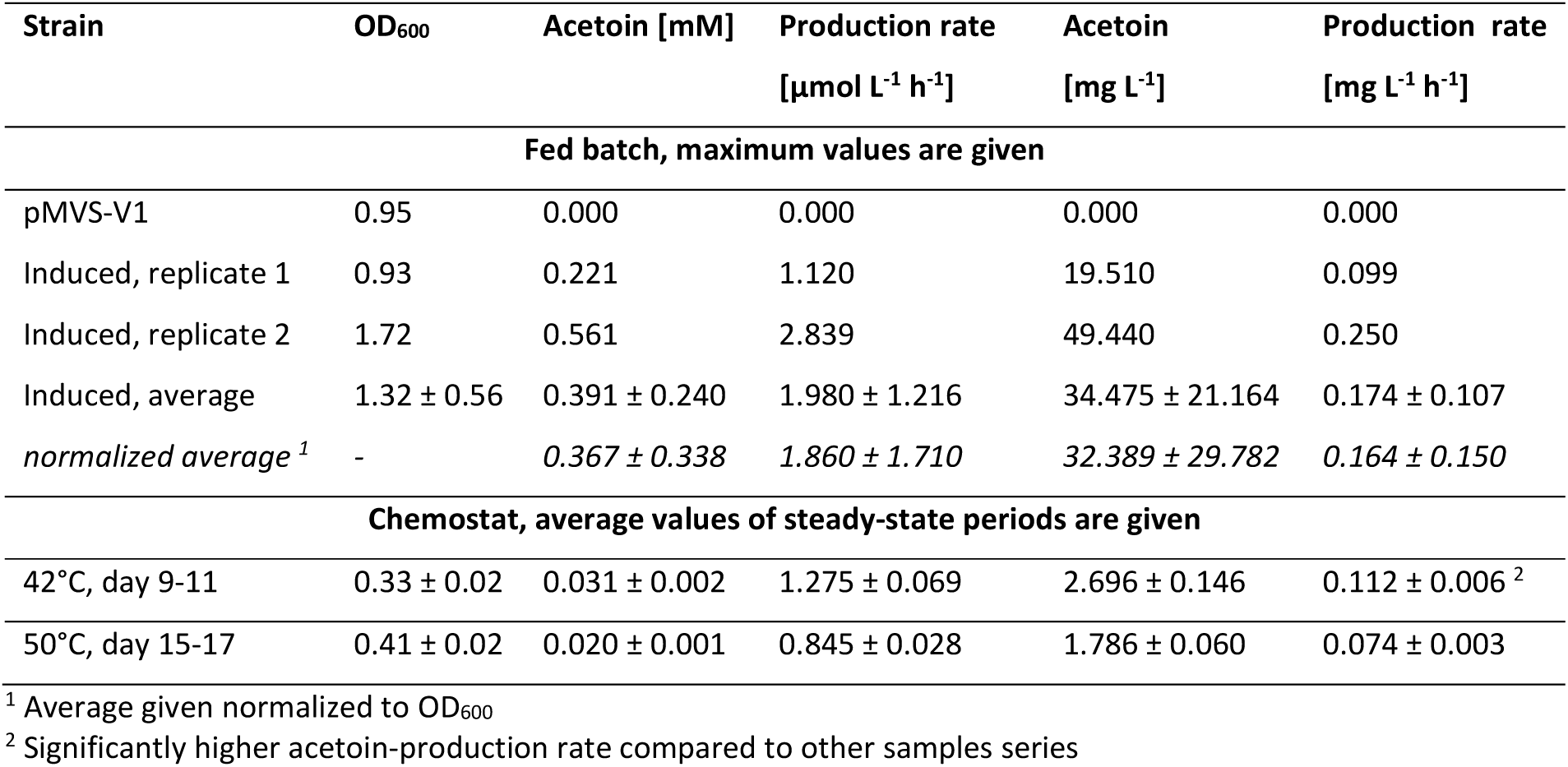
Summary of growth and acetoin productivity across the different bioreactor experiments.

Finally, based on our observations, we hypothesized that acetoin production should be possible in a continuous system, representing a situation that is closer to an actual bioprocess. Thus, we operated a chemostat with the *M. thermautotrophicus* [pMVS1111a_P*_tet_*_alsSD] strain (**Figure 5**). Again, we operated the bioreactors at 42°C and we induced acetoin production by continuous addition of aTc. Shortly after induction, we observed a peak in acetoin production on day 8 at a volumetric production rate of 1.80 µmol L^-1^ h^-1^ (titer, 0.043 mM) which stabilized at a rate of 1.28 ± 0.07 µmol L^-1^ h^-1^ (titer, 0.031 ± 0.002 mM) (**Figure 5**, **Table 4**). This represents a similar volumetric production rate compared to the (fed-)batch experiments before. However, the specific production rate (normalized to OD_600_) was approximately two-fold higher compared to the fed-batch bioreactor (**Table 5**). To investigate the effect of the growth temperature on acetoin production, we then increased the temperature in the bioreactor to 50°C, which resulted in a stable but significantly (p=0.000029) reduced volumetric acetoin production rate of 0.84 ± 0.03 µmol L^-1^ h^-1^ (0.020 ± 0.001 mM) (**Figure 5**, **Table 4**). Further increasing the temperature to 60°C led to an increase in OD_600_, while we did not measure any acetoin anymore (**Figure 5**). The increased growth was accompanied by an increase in substrate gas consumption and CH_4_ production rates (**Supplemental Figure 3**).

**Figure 5.**
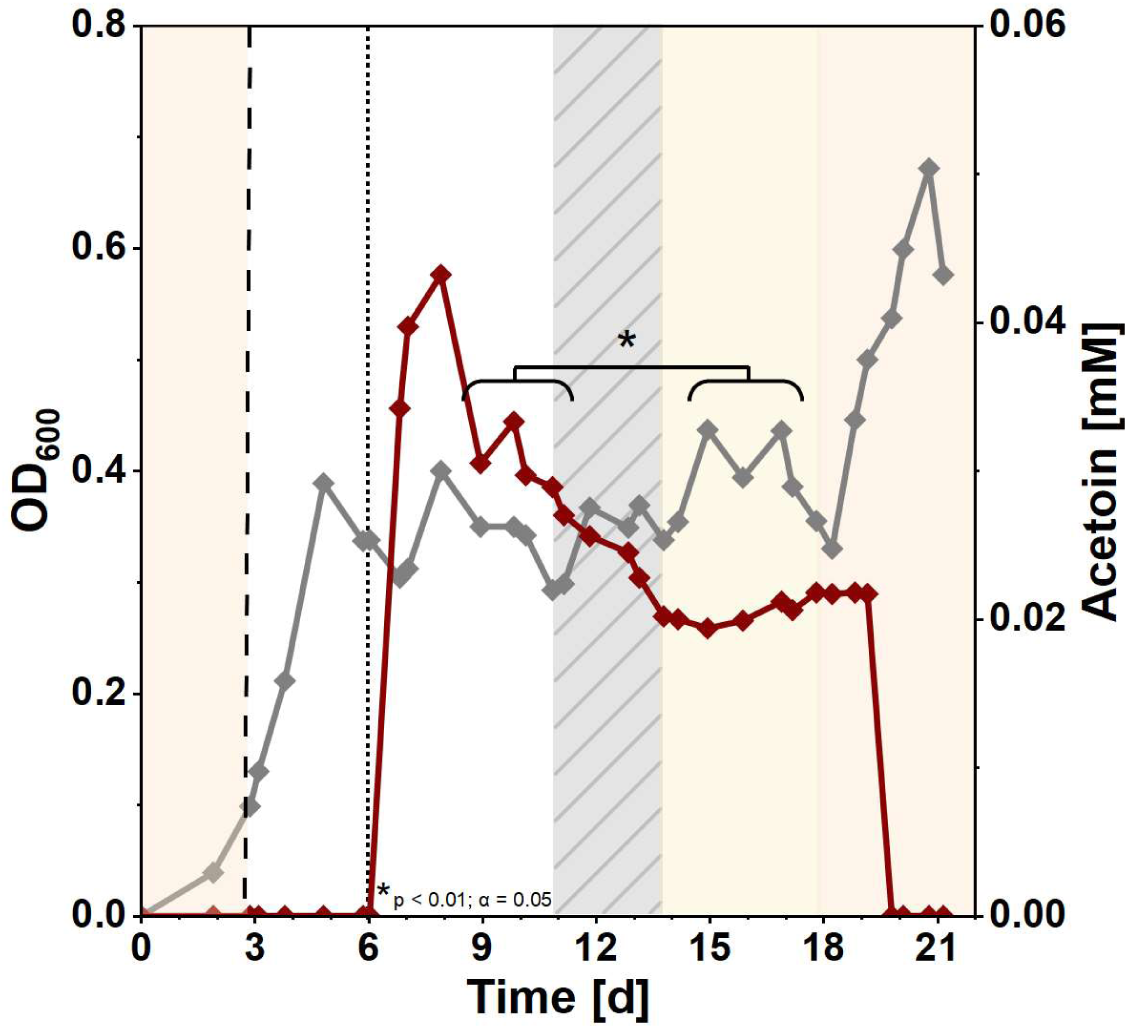
Recombinant acetoin production with *M. thermautotrophicus* [pMVS1111a_P*_tet_*_alsSD] in a chemostat bioreactor. Growth (OD_600_, grey) and acetoin production (red). Acetoin production was induced with 1 µg mL^-1^ anhydrotetracycline (dotted line) after approximately 6 days. Light orange shading, cultivation temperature set to 60°C, light yellow shading, cultivation temperature set to 50°C; dashed line, switch from batch to continuous mode and decrease of cultivation temperature to 42°C; grey-hatched area, technical disturbances, n=1. The asterisk indicates significantly different production rates between the two periods.

**Table 5.**
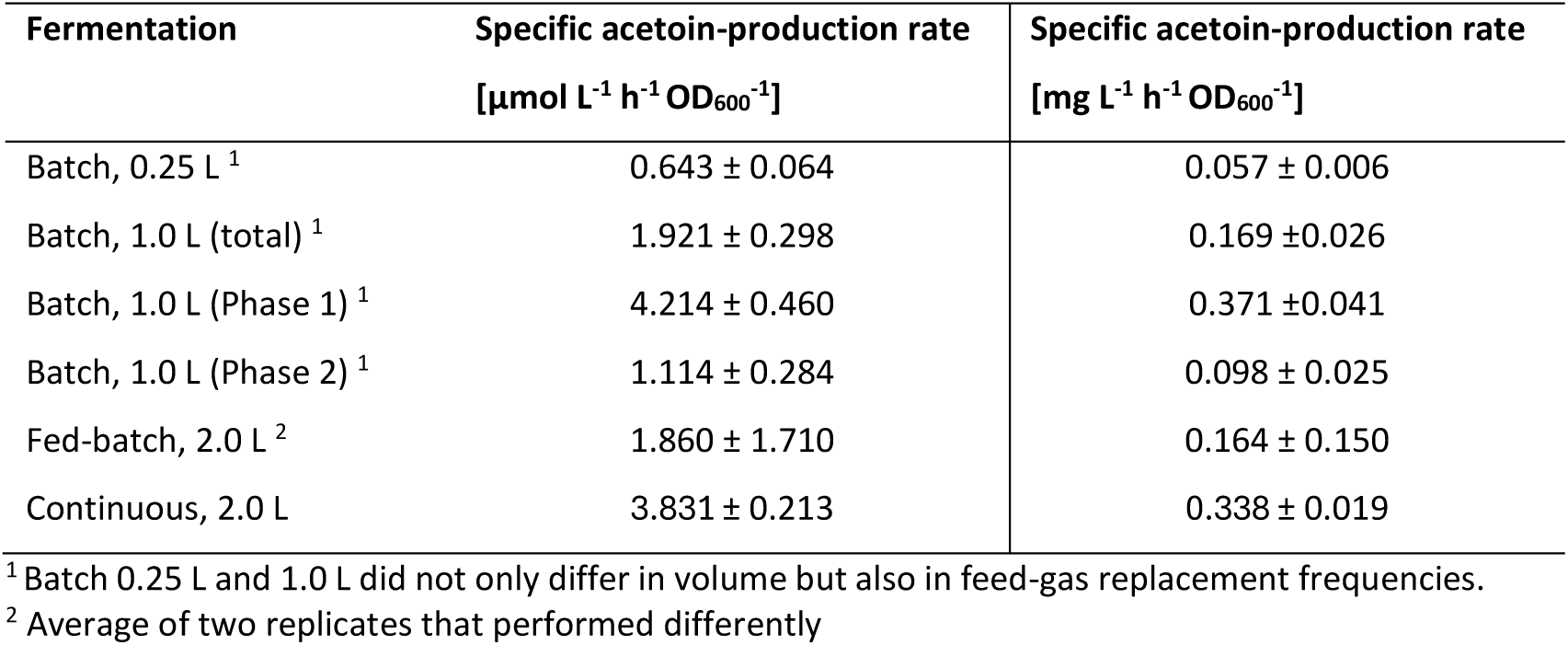
Comparison of specific acetoin-production rate during induced state in differing cultivation vessels.

| Fermentation | Specific acetoin-production rate<br>[μmol L<sup>-1</sup> h<sup>-1</sup> OD<sub>600</sub><sup>-1</sup>] | Specific acetoin-production rate<br>[mg L<sup>-1</sup> h<sup>-1</sup> OD<sub>600</sub><sup>-1</sup>] |
| --- | --- | --- |
| Batch, 0.25 L <sup>1</sup> | 0.643 ± 0.064 | 0.057 ± 0.006 |
| Batch, 1.0 L (total) <sup>1</sup> | 1.921 ± 0.298 | 0.169 ± 0.026 |
| Batch, 1.0 L (Phase 1) <sup>1</sup> | 4.214 ± 0.460 | 0.371 ± 0.041 |
| Batch, 1.0 L (Phase 2) <sup>1</sup> | 1.114 ± 0.284 | 0.098 ± 0.025 |
| Fed-batch, 2.0 L <sup>2</sup> | 1.860 ± 1.710 | 0.164 ± 0.150 |
| Continuous, 2.0 L | 3.831 ± 0.213 | 0.338 ± 0.019 |

<sup>1</sup> Batch 0.25 L and 1.0 L did not only differ in volume but also in feed-gas replacement frequencies.
<sup>2</sup> Average of two replicates that performed differently

## 4. Discussion

We demonstrated the successful implementation of an aTc-inducible promoter system, which allowed the completely autotrophic recombinant production of acetoin in *M. thermautotrophicus* from H_2_ and CO_2_ in a defined mineral salt medium in batch bottles, fed-batch- and chemostat bioreactors. The induction of the expression of a thermostable β-galactosidase that we had previously used to study the strength of various constitutive promoters (Fink et al., 2021) was optimal at an aTc concentration of 1 µg mL^-1^. We observed a basal expression of the uninduced system, which was higher than for the empty vector control, potentially due to leaky expression of the system. However, experimental variation, such as inconsistent temperatures within the incubator due to sampling events, might have an impact on this observation. Important to note is that we used 60°C for those experiments, because the temperature-dependency of the TetR-binding to the operator was demonstrated recently for *Geobacillus stearothermophilus* with derepression at temperatures above 60°C (Mol, 2022). In fact, this temperature induction was now implemented for *M. marburgensis*, resulting in repression of gene expression at 55°C and induction of gene expression at 62°C (Unger et al., 2025). A basal activity was also observed in *M. marburgensis* at 55°C (Unger et al., 2025). Temperature fluctuations or uneven temperature distribution within the incubator may have caused aTc-independent induction in our experiments, which may have increased the experimental variation. Nevertheless, we observed aTc induction of ∼3.7-fold compared to the uninduced control. For the induction of acetoin production, this effect should be less relevant because we operated at a temperature of 42°C to minimize risks of inactivation of the recombinant enzymes from *S. thermophilus*, which has an optimum growth temperature of 42°C (Radke-Mitchell and Sandine, 1986). In fact, we confirmed the functionality of the aTc-inducible system because no considerable recombinant acetoin production was observed without aTc induction during growth.

The non-specific accumulation of acetoin in all strains, including the wildtype, empty vector, and uninduced control strains, was most likely due to accumulation of the intermediate acetolactate, which is natively produced by *M. thermautotrophicus* in valine biosynthesis from pyruvate (Kaster et al., 2011; Mühling et al., 2024; Unger et al., 2025). We speculate that during growth amino acids are immediately incorporated into biomass, whereas when growth ceases intermediates may accumulate. Acetolactate can then chemically decompose into acetoin and CO_2_ without enzymatic catalysis. Potentially, the decomposition of biomass provides more precursors that could increase this effect. Interestingly, the acetolactate decarboxylase (AlsD) from *S. thermophilus* was found to be activated by the presence of branched-chain amino acids, leading to higher activity (Monnet et al., 2003). It has been shown in recent years that methanogens actively excrete amino acids, potentially as part of an overflow metabolism to balance surplus electron carriers (Taubner et al., 2023). Such a situation would be especially relevant when H_2_ and CO_2_ are available in excess while no flux is directed into biomass production. While the exact mechanisms of such overflow metabolism are not understood, our results show that acetoin only non-specifically accumulates when growth is no longer observed in batch experiments and does not at all accumulate in the fed-batch bioreactor, and thus are in line with this scenario.

In chemostats, we achieved the highest acetoin productivity at a temperature of 42°C. Importantly, at 50°C we achieved slightly reduced acetoin production rates. This further indicates suboptimal enzyme conditions, because at the same time the growth (OD_600_) and CH_4_ production rates increased, as expected at a more optimal temperature for *M. thermautotrophicus*. However, a changed flux distribution within the cell without reduced activities of the recombinant enzymes can also be a reason for this observation. When elevating the temperature to 60°C, growth and CH_4_ production further surged, while acetoin was not detectable anymore.

It is unlikely that the increased temperature had a negative impact on the induction of the aTc-inducible system, because we established the system at 60°C and did not observe considerable unspecific temperature-induced expression. In fact, a temperature-induced increase in gene expression (Mol, 2022; Unger et al., 2025) would have rather elevated the enzyme levels and led to higher acetoin bioproduction, in case of stable enzyme activity. Instead, we speculate that the halt of the acetoin production is due to inactivation/denaturation of the enzymes at elevated temperatures. However, we cannot fully rule out that acetoin is stripped out of the bioreactor with the off-gas, although acetoin has a boiling point of ∼148°C. Gas stripping could also be a factor that reduced the apparent concentration of acetoin already at 42°C and 50°C and needs to be experimentally tested in the future.

Plasmid-based gene expression offers benefits for laboratory-scale testing of production pathways. However, to avoid the need for antibiotic selection and effects, such as variation of plasmid-copy numbers, further optimization of the system should be performed with genome-integrated acetoin-production operons, which is possible with the available genetic tools (Fink et al., 2023; Klein et al., 2025). We did not further investigate whether regulation of native enzymes, recombinant enzyme activities under sub-optimal conditions, unbalanced expression levels of *alsS* and *alsD*, allosteric inhibition, or other factors reduced the flux toward acetoin. Engineering the regulation of pathway enzymes will have to be considered for improving productivity. One strategy is enzyme engineering of the native enzymes, because alleviation of allosteric inhibition of the acetolactate synthetase from *M. thermautotrophicus* and recombinant production in *M. marburgensis* could be implemented to increase the flux toward valine bioproduction in which acetolactate is an intermediate (Unger et al., 2025). Engineering the dependence of the presence of branched-chain amino acids for AlsD activity from *S. thermophilus* (Monnet et al., 2003) could be the starting point of further optimization of the flux toward acetoin. In addition, identifying more temperature-stable enzyme variants should be considered (Jia et al., 2017; Kato et al., 2024).

## 5. Conclusion

Recombinant bioproduction with thermophilic methanogens can lead to more versatile and economically viable power-to-x processes that can be integrated with existing infrastructure. Our results demonstrate that this strategy is possible in practice, exemplified with recombinant acetoin bioproduction. However, an increase in productivity will be central to make this platform applicable for biotechnological purposes. In combination with the development of further genetic tools for Methanobacteriales (Klein et al., 2025; Unger et al., 2025), the development of strategies to implement relevant and achievable bioproduction routes (Contreras et al., 2022; Mühling et al., 2024), and hypothesis testing with genome-scale metabolic modeling (Casini et al., 2023), we will now be able to move this technology one step further beyond biomethanation.

## Author contributions

Tina Baur (T.B.), Bastian Molitor (B.M.), and Largus T. Angenent (L.T.A.) designed the experiments. T.B. and Gabriela Contreras (G.C.) performed the genetic work. T.B. characterized the inducible promoter and performed the acetoin production experiments in bottles. Maximilliene T. Allaart (M.T.A.) and Aaron Zipperle (A.Z.) planned and performed the bioreactor experiments and analyzed the data together with T.B. and B.M.. Statistical analyses were done by Christian Fink (C.F.). B.M. and L.T.A. supervised the work and acquired funding. T.B., M.T.A., C.F., and B.M. prepared figures and tables. T.B., M.A., G.C., and A.Z. drafted the Materials and Methods, B.M. drafted the remaining manuscript, and all authors contributed to editing and approved the final version.

## Acknowledgments

The authors thank Dr. Sebastian Beblawy for designing the codon-optimized acetoin-production operon, Kurt Gemeinhardt and Dr. Daniel Buchner for support with GC-MS analyses, and Dr. Andrés Ortiz-Ardila for support with statistical analyses.

