## Supplementary Material for "Metabolic engineering of *Methanothermobacter thermautotrophicus* ΔH for recombinant acetoin production"

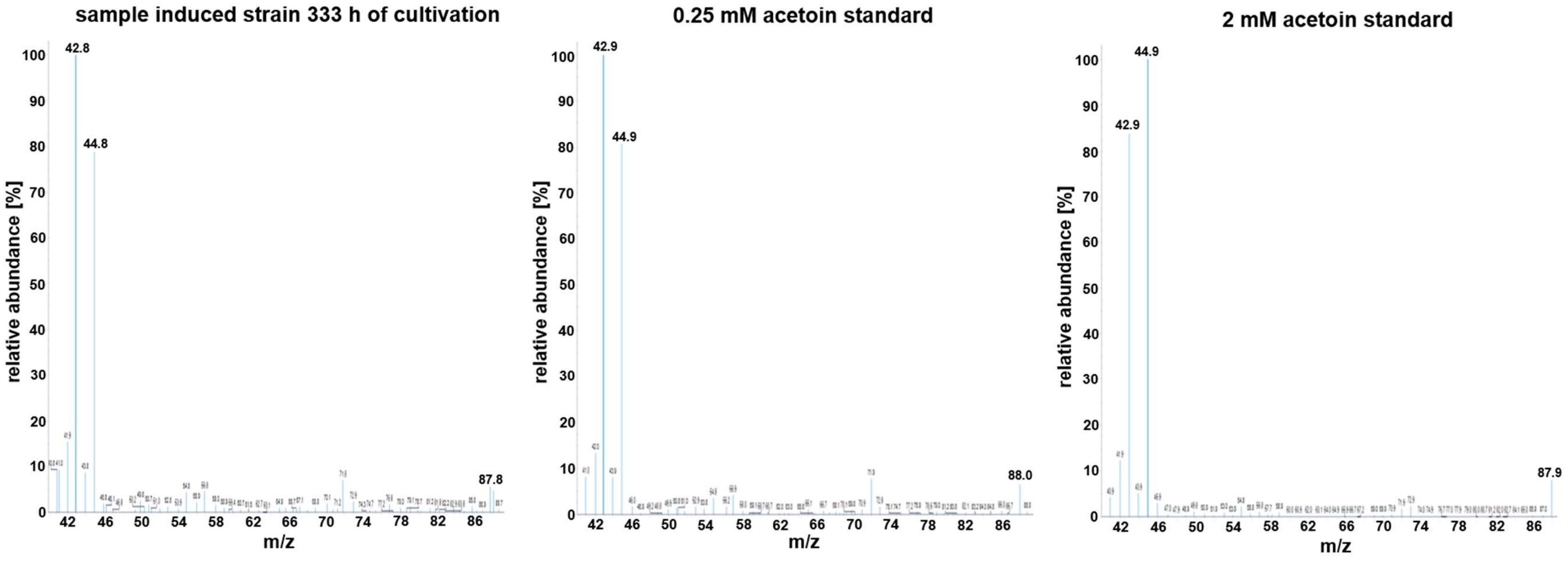


**Supplemental Figure 1. Mass spectra of chromatographic acetoin peaks of analyzed samples.** Left, representative sample of induced *M. thermautotrophicus* [pMVS1111a_$P_{tet}$_alsSD] culture (after 333 h of cultivation); middle, defined 0.25 mM acetoin standard; right, defined 2 mM acetoin standard. A comparison of the spectra with the NIST database confirmed the identity of the compound as acetoin.

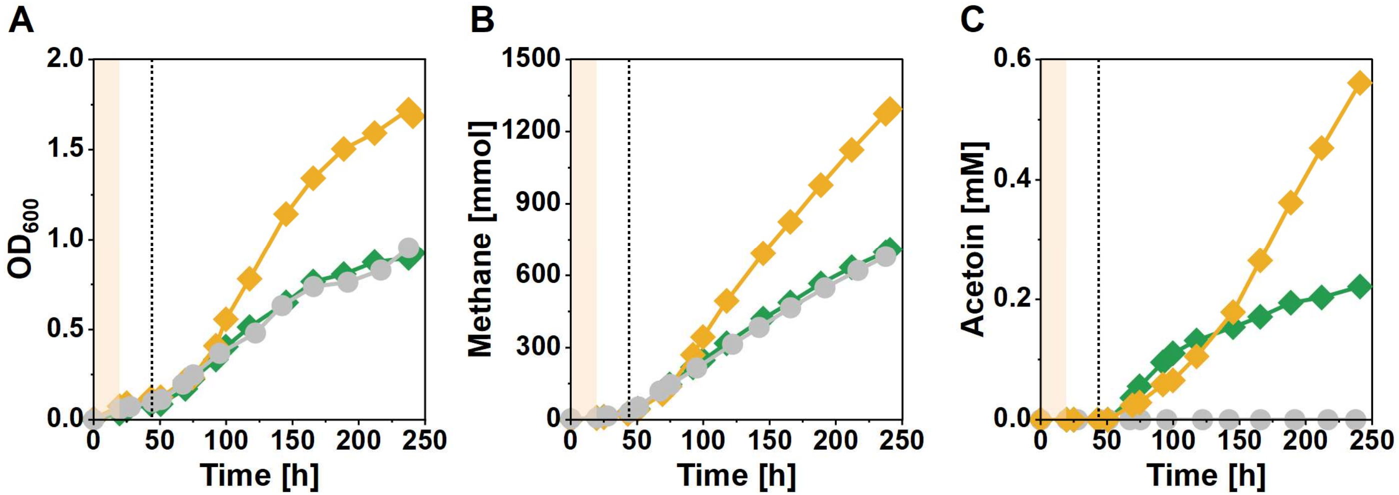


**Supplemental Figure 2. Recombinant acetoin production with *M. thermautotrophicus* in fed-batch bioreactors. A**, Growth ($OD_{600}$); **B**, methane production; **C**, acetoin production. Grey circles, *M. thermautotrophicus* [pMVS-V1] (n=1); green diamonds, *M. thermautotrophicus* [pMVS1111a_P*tet*_alsSD] replicate 1; yellow diamonds, *M. thermautotrophicus* [pMVS1111a_P*tet*_alsSD] replicate 2. Initial growth occurred at 60 °C (light orange shading), afterwards, the temperature was decreased to 42 °C and acetoin production was induced using 1 µg $mL^{-1}$ anhydrotetracycline (dotted line).

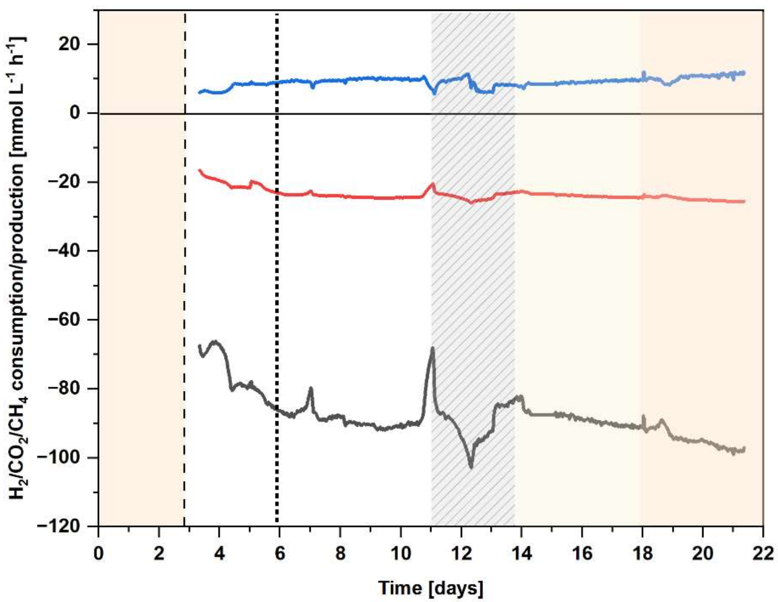


**Supplemental Figure 3. Gas data from chemostat bioreactor experiment related to Figure 5 in main text**. 5-point moving average of consumption- (grey, $H_2$ and red, $CO_2$) or production- (blue, $CH_4$) rates are shown. For reference, acetoin production was induced with 1 µg $mL^{-1}$ anhydrotetracycline (dotted line) after approximately 6 days. Light orange shading, cultivation temperature set to 60°C, light yellow shading, cultivation temperature set to 50°C; dashed line, switch from batch to continuous mode and decrease of cultivation temperature to 42°C; grey-hatched area, technical disturbances. n=1.
